# Good tutors are not Dear Enemies in Song Sparrows

**DOI:** 10.1101/112516

**Authors:** Çağlar Akçay, S. Elizabeth Campbell, Michael D. Beecher

## Abstract

Bird song is the most widely studied example of vocal learning outside human language and shares important parallels with it, including the importance of social factors during development. Our understanding of how social factors affect song learning however remains surprisingly incomplete. Here we examine the possible role of aggressive interactions in determining song “tutor” choice in song sparrows (*Melospiza melodia*), a songbird in which individuals display song learning strategies ranging from learning primarily from one tutor, to learning a few songs each from a number of tutors. We test two hypotheses: The Competition hypothesis suggests that young birds learn more from tutors with whom they compete especially intensely and predicts that tutees will respond with high aggression to tutor songs. In contrast the Cooperation hypothesis suggests that song learning reflects a cooperative relationship between the tutor and the tutee and predicts that tutees will respond with low aggression to tutor songs. In a playback experiment we found that birds respond more aggressively to songs of their tutors than they do to songs of strangers and that the strength of aggressive response correlated positively with how much they had learned from that tutor. These results provide the first field evidence for the hypothesis that young males preferentially learn their songs from adult males with whom they compete most intensely during the song-learning phase, and perhaps afterwards.

## Introduction

Although vocal communication is ubiquitous in the animal kingdom, social learning of vocal signals is limited to a few taxa, including humans (but not other primates), cetaceans, bats, elephants and three orders of birds (Baptista & Schuchmann, 1990; Boughman, 1998; Marler & Tamura, 1964; Pepperberg, 1994; Reiss & McCowan, 1993). Of these, song in songbirds is the best studied system next to human language (Beecher & Brenowitz, 2005; Catchpole & Slater, 2008).

Early studies showed striking parallels between the development of vocal signals in humans and songbirds including an early sensitive period, a predisposition to learn conspecific vocalizations, a babbling (or subsong) stage, and the necessity of auditory feedback for normal development (Marler, 1970). Another parallel is the social nature of vocal development. Although the social aspect of vocal development is obvious for humans, the potent role of social interactions in song learning was not fully appreciated until laboratory studies used live birds as song “tutors” rather than recorded song as had been conventional (Baptista & Petrinovich, 1984, 1986). Although it is now widely accepted that song learning is a social process in which young birds (tutees) hear and engage in interactions with adults (tutors), there is a dearth of studies attempting to identify the critical social factors (Beecher, 2008).

Despite the many striking parallels between human and songbird vocal learning, the key social factors in vocal learning may be quite different for the two taxa. In particular, whereas the tutor-tutee (teacher-student) relationship in humans is clearly a cooperative one, with both parties typically related, the common case in songbirds is that tutor and tutee are unrelated competitors. This is because most songbirds commence song-learning only after they disperse from their natal area, and thus their song tutors are future territorial competitors rather than their parents usually fathers (here and throughout the rest of paper we focus on male song and use the male pronoun, but note that female singing is common in other species, particularly in the tropics; Riebel, Hall, & Langmore, 2005). Species where fathers act as song tutors for their sons are rare (Grant & Grant, 1996; Greig, Taft, & Pruett-Jones, 2012; Immelmann, 1969). Instead, in most songbirds song tutors are unrelated adults some of whom will be their territorial competitors come the next breeding season (Beecher & Brenowitz, 2005; Brenowitz & Beecher, 2005). This point can be illustrated with song sparrows living in Washington State (Beecher, 2017). In this population, song learning occurs in the period after natal dispersal and before the bird’s first breeding season the following spring. Neighbors typically ‘share’ song types, and this song sharing has been shown to be a result of song learning (Beecher, 2008; Beecher, Campbell, & Stoddard, 1994; Nordby, Campbell, & Beecher, 1999). The period of song learning also coincides with territory establishment during which young birds also engage in aggressive interactions with their future neighbors (Arcese, 1989; Nice, 1943), and shared songs are used by adult birds as part of a graded signaling system in aggressive interactions (Akçay, Tom, Campbell, & Beecher, 2013; Burt, Campbell, & Beecher, 2001). All of these lines of evidence suggest that song learning may be influenced by the amount of aggressive and competitive interactions between the tutors and the young birds. We term this hypothesis the *“Competition”* hypothesis.

A different line of thinking, however, suggests that even under these circumstances the songbird tutor-tutee relationship could be an at least partially cooperative one. As has been shown for numerous diverse taxa, territorial neighbors often enter into a ‘Dear Enemy’ relationship where they are more tolerant of their neighbors than they are of strangers (Akçay et al., 2009; Fisher, 1954; Temeles, 1994). Hence it is possible that it might actually benefit an established territorial adult to ‘teach’ his songs to a young bird who is in a position to become his future neighbor; in short, a ‘dear tutor-tutee’ relationship could underlie and support a ‘dear enemy’ relationship. This idea can be seen as an extension of the observation that that in many group-living species with vocal learning (e.g. dolphins, parrots and cooperatively breeding songbirds), vocal learning seems to have an affiliative function in which individuals learn their vocalizations from members of their social groups (Akçay, Hambury, Arnold, Nevins, & Dickinson, 2014; Berg, Delgado, Cortopassi, Beissinger, & Bradbury, 2012; Brown, Farabaugh, & Veltman, 1988; Price, 1998; Sharp, McGowan, Wood, & Hatchwell, 2005). This *“Cooperation”* hypothesis is also consistent with recent reviews of animal ‘teaching’ in which the putative tutor does not obtain immediate benefits (and may even pay immediate costs) by engaging in the teaching of a tutee (Hoppitt et al., 2008). Under this view, older birds would reap some form of delayed benefit from tutoring young birds and having them as neighbors. This benefit might come in the form of increased breeding success (Beletsky & Orians, 1989), access to extra-pair females that are mated to the young males or less competition for within-pair paternity from these young males (Hill, Akçay, Campbell, & Beecher, 2011). The last two potential benefits are based upon the findings that in many songbirds including song sparrows the older males are more likely to successfully pursue extra-pair matings (Akçay et al., 2012; Hill et al., 2011; Hsu, Schroeder, Winney, Burke, & Nakagawa, 2015).

Here we present a test of competition and cooperation hypotheses in song sparrows. Song sparrows are close-ended learners who learn their songs in the period after dispersal from the natal area and the beginning of their first breeding season the following spring and do not change their song repertoire in subsequent years (Nordby, Campbell, & Beecher, 2002). Extensive field studies have shown that while on average a bird copies about half of his 8 or 9 songs from a single tutor (the best tutor) and the rest from multiple other tutors, there is a range of learning strategies, varying from copying all his songs from a single tutor to copying a single song from each of 8 or 9 tutors (Akçay, Campbell, Reed, & Beecher, 2014; Beecher et al., 1994; Nordby et al., 1999; Nordby, Campbell, & Beecher, 2007). As noted above, adult song sparrows use shared songs as part of a graded signaling system that may indicate the primary role of competitive interactions in song learning. At the same time, multiple studies have shown that male song sparrows can individually recognize their neighbors and generally show reduced aggression to neighbors compared to strangers (Akçay, Reed, Campbell, Templeton, & Beecher, 2010; Akçay et al., 2009; Wilson & Vehrencamp, 2001), suggesting the opportunity for cooperative interactions with potential tutors exists.

We tested the two hypotheses by asking whether a bird would be respond more or less aggressively to a simulated intrusion by a former tutor compared to a stranger, and whether aggressive response would vary with how much the young bird had learned from the tutor. Specifically, in a playback experiment to subjects with known song learning histories, we compared their aggressive response to the tutor from whom they had learned the most, to their aggressive response to songs from a stranger. The cooperation hypothesis predicts that tutees should respond *less* aggressively to their best tutors than to strangers, and less aggressively to tutors from whom they learned more than from tutors from whom they learned less. In contrast, the competition hypothesis predicts precisely the opposite: subjects should respond *more* aggressively to their best tutors than to strangers, and more aggressively to tutors from whom they learned more than from tutors from whom they learned less.

## Methods

### (a) Study site and subjects

We studied a banded population of song sparrows in Discovery Park, Seattle, Washington, USA. Between 2009 and 2014 all the territorial males (about ~120 males each year) were banded with a US Fish and Wildlife Service metal band and three colored bands. As a part of our long term study on song learning (Beecher, 2008), the complete song repertoire of each male was also recorded with Marantz PMD 660 recorders and Sennheiser ME66/K6 shot-gun microphones. The full repertoire was considered to be recorded after at least 16 song switches (Nordby et al., 1999). Subjects in the playback experiment were 13 banded and recorded male song sparrows in our study population in Discovery Park, Seattle, Washington, USA. We tested each subject twice on different days with a counterbalanced order for two trial types. The experiments were conducted from March 18 to April 14, 2014.

### (b) Tracing song learning

We chose males with known ages and song learning histories that held territories in Spring 2014. Three of the subjects hatched in 2009, 6 in 2010 and 7 in 2011. All the subjects were banded either in juvenile plumage (before their first molt) or singing plastic song before their first Spring when the songs crystallize (around March 1^st^). We made sonagrams of all the songs in the repertoire of the tutees and potential tutors using Syrinx (www.syrinxpc.com, John Burt, Seattle, WA). We printed out several variations of each song of all the males. The tutors for each tutee were determined as described in detail in our previous studies (Akçay, Campbell, Reed, et al., 2014; Nordby et al., 1999). Three judges visually compared the songs of the tutees and tutors independently and laid out matching songs on a large table. After this step, the three judges discussed their best match decisions, and arrived at a consensus. If a single adult had the best matching song for a given tutee song, that tutor got a credit of 1 for that song. If more than one adult had equivalently good matches for a given tutee song, then each tutor got credit of 1/N, where N was the number of tutors with equally good matching songs. Because of the high level of song sharing in our population (Hill, Campbell, Nordby, Burt, & Beecher, 1999) splitting credit between multiple tutors happens about half the time (Akçay, Campbell, Reed, et al., 2014).

### (c) Design and stimuli

Subjects were tested with two songs each from 1) the male with the highest tutoring score for that bird (the bird’s ‘best’ tutor) and 2) a stranger male that held a territory at 1 to 2 km from the territory of the subject. In all cases the best tutor was no longer present in the study area, most likely to due to death as territorial males do not make significant moves (Akçay, Campbell, & Beecher, 2015; Arcese, 1989). In seven out of 13 cases, the tutor had been a neighbor with the subject. In the remaining six cases, the tutor had not shared an immediate boundary with the subject but was within two territories of the subject’s territory. Previous research in other songbirds has shown that males remember and recognize their familiar neighbors even after these disappear (Godard, 1991; McGregor & Avery, 1986). We therefore expected that subjects would be able to recognize their tutors.

We carried out the playbacks at the center of the subjects’ territories to have a standardized location for contrasting responses to strangers vs. tutors. Previous studies in song sparrows (and most of the other songbirds studied) have shown no difference in response strength between stranger playback and a randomly chosen familiar bird, usually a neighbor, at the territory center (Stoddard, 1996) We reasoned therefore that getting a difference in response strength to tutors compared to strangers in either direction would be stronger test of the alternative hypotheses. We note also that since the tutors had disappeared by the time of the playbacks, and some of the best tutors did not share a boundary with the subject to start with, there was no current shared boundary between subjects and best tutors.

The stimuli for the best tutors were chosen from the songs they shared with the tutee (i.e. from the songs that the tutee had learned from the tutor). Stranger songs were non-shared with the subject (as stranger songs almost always are). Playback tapes were created in Syrinx so that stimulus songs (a single rendition per song type) would be presented every ten seconds.

### (d) Playback procedure

Each subject was tested twice, once with the songs of his best tutor and once with the songs of a stranger on different days not farther apart than 1 week. The order was counterbalanced across subjects. We started each trial by setting up a speaker (iMainGo, Portable Sound Laboratories, Inc) at the center of the subject’s territory. The speaker was connected to an iPod with a 20 m cable. The stimuli were played at approximately 80 dB SPL, measured at 1 m (Radio Shack 33-2055 sound meter), corresponding to normal broadcast song amplitude. Two observers recorded the behavior of the subject (flights, songs, wing waves, distance with every flight) verbally using the same recording equipment as above. Three minutes after the first sighting of the subject, we switched to the second song type and carried on the trial for another three minutes.

### (e) Response measures and data analyses

From the trial recordings we extracted the following response measures: duration of the trial (from first sighting of the male to the last playback), number of flights, time spent within 5m of speaker, closest approach to the speaker. The numbers of flights and songs were converted to rates per minute to account for unequal duration of observation across trials due to different latencies to respond.

We use rate of flights, proportion of trial spent within 5m and closest approach distance as our primary variables of aggression. As these variables were highly correlated with each other we used a principal component analysis (PCA, unrotated, correlation matrix) to arrive at a single aggression score (see the correlation matrix in Table 1). Our previous studies with taxidermic mounts indicate that aggression scores calculated from these measures reliably predict attack (Akçay, Campbell, & Beecher, 2014; Akçay et al., 2013). These three variables also constitute behaviors that are expression of a “behavioral character” (Araya-Ajoy & Dingemanse, 2014) corresponding to aggressiveness in this species (Akçay et al., 2015). The first component of the PCA explained 73.6% of the variance and we took these scores as the aggression scores (see Table 1 for coefficients). Higher aggression scores meant higher levels of aggressive response. We then ran a mixed ANOVA on the aggression scores with the condition as a within-subject factor and proportion learned from best tutor as a between subject covariate.

**Table 1.**
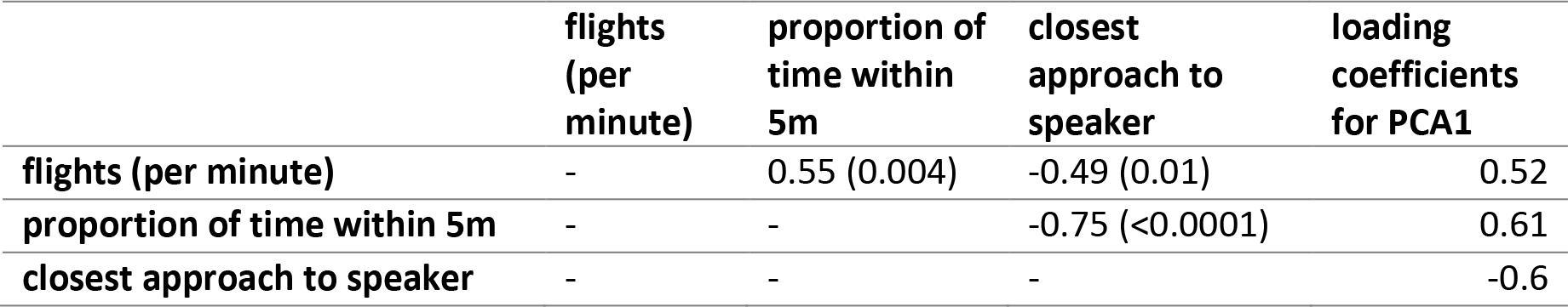
Correlation coefficients (p-values) between aggressive behaviors and the loading coefficients on the PCA (rightmost column).

|  | **flights**<br>**(per**<br>**minute)** | **proportion of**<br>**time within**<br>**5m** | **closest**<br>**approach to**<br>**speaker** | **loading**<br>**coefficients**<br>**for PCA1** |
| --- | --- | --- | --- | --- |
| **flights (per minute)** | - | 0.55 (0.004) | -0.49 (0.01) | 0.52 |
| **proportion of time within 5m** | - | - | -0.75 (<0.0001) | 0.61 |
| **closest approach to speaker** | - | - | - | -0.6 |

### (f) Ethical statement

This research was conducted in accordance with the ABS/ASAB Guidelines for the treatment of animals in behavioural research and teaching, and with approval from the University of Washington Institutional Animal Care and Use Committee (IACUC # 2207-03) and the U.S. Fish and Wildlife Service banding permit (banding permit # 20220). We did not observe any adverse effect of handling and banding the birds in our long term study as banding time is minimized and birds returned to their territories immediately (within minutes of capture). All subjects were banded at least one year before the experiment and had been holding their territories for at least one year before the experiment.

## Results

Repertoire sizes of subjects ranged from 6 to 12 song types with a mean of 9.15. The number of tutors for each tutee ranged from 1 to 7. On average, the best tutors accounted for 48% of the songs in the repertoires of the tutees (range: 15% to 83.3%).

Subjects responded more strongly to tutor playback than to stranger playback (F_1,11_= 13.61, p= 0.004, Figure 1). 12 out of 13 subjects for whom we had both trials responded more aggressively to the tutor than to the stranger. There was no main effect of proportion of the song repertoire learned from the best tutor (F_1,11_= 0.03, p=0.88) but there was a significant interaction between proportion of song repertoire learned from best tutor and condition (i.e. tutor vs. stranger), F_1,11_= 5.93, p= 0.03. The more the tutee had learned from the best tutor, the more aggressive was his response to that bird’s song compared to his response to the stranger song (Figure 2).

**Figure 1.**
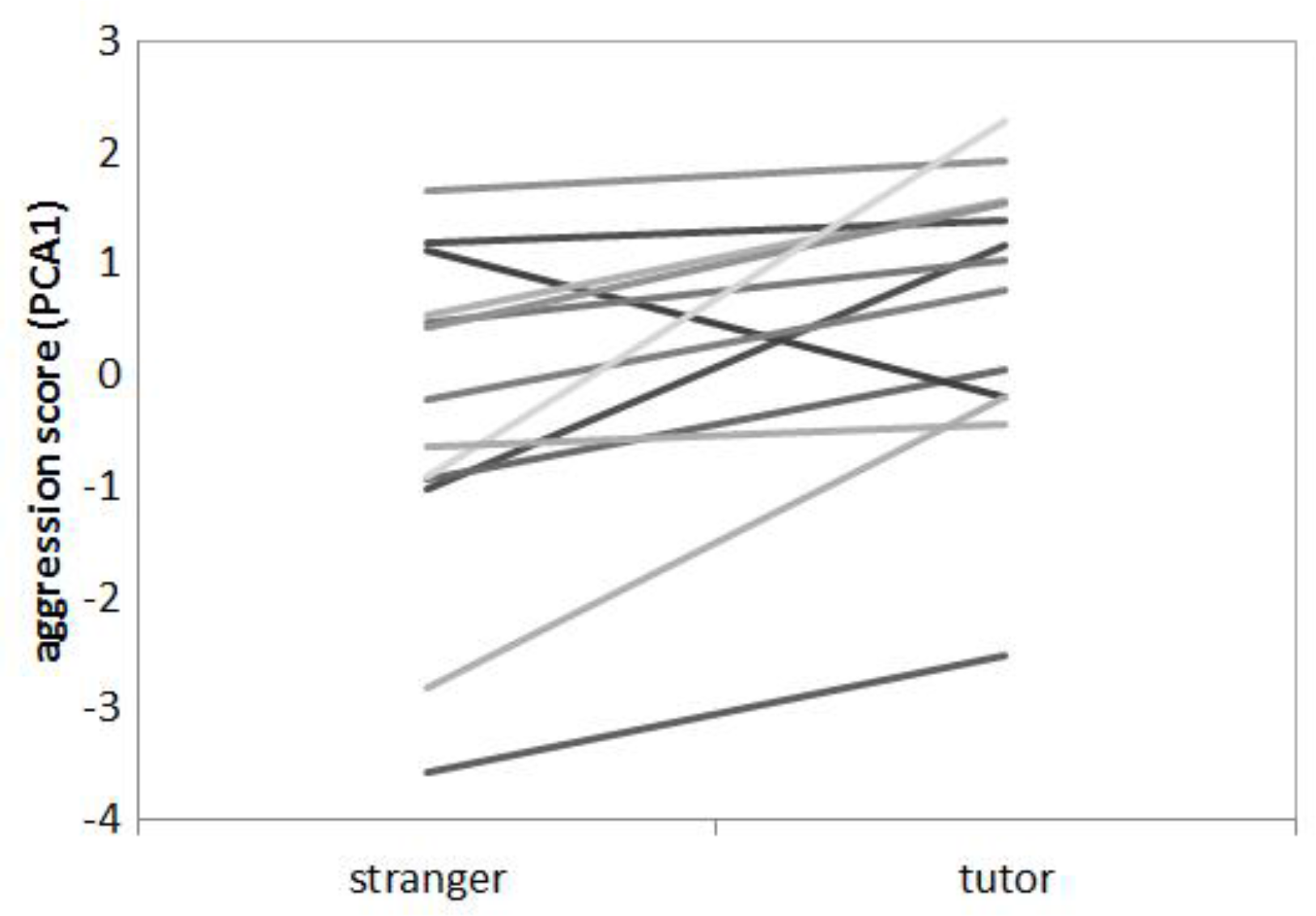
Aggression scores in stranger vs. tutor trials for individual subjects.

**Figure 2.**
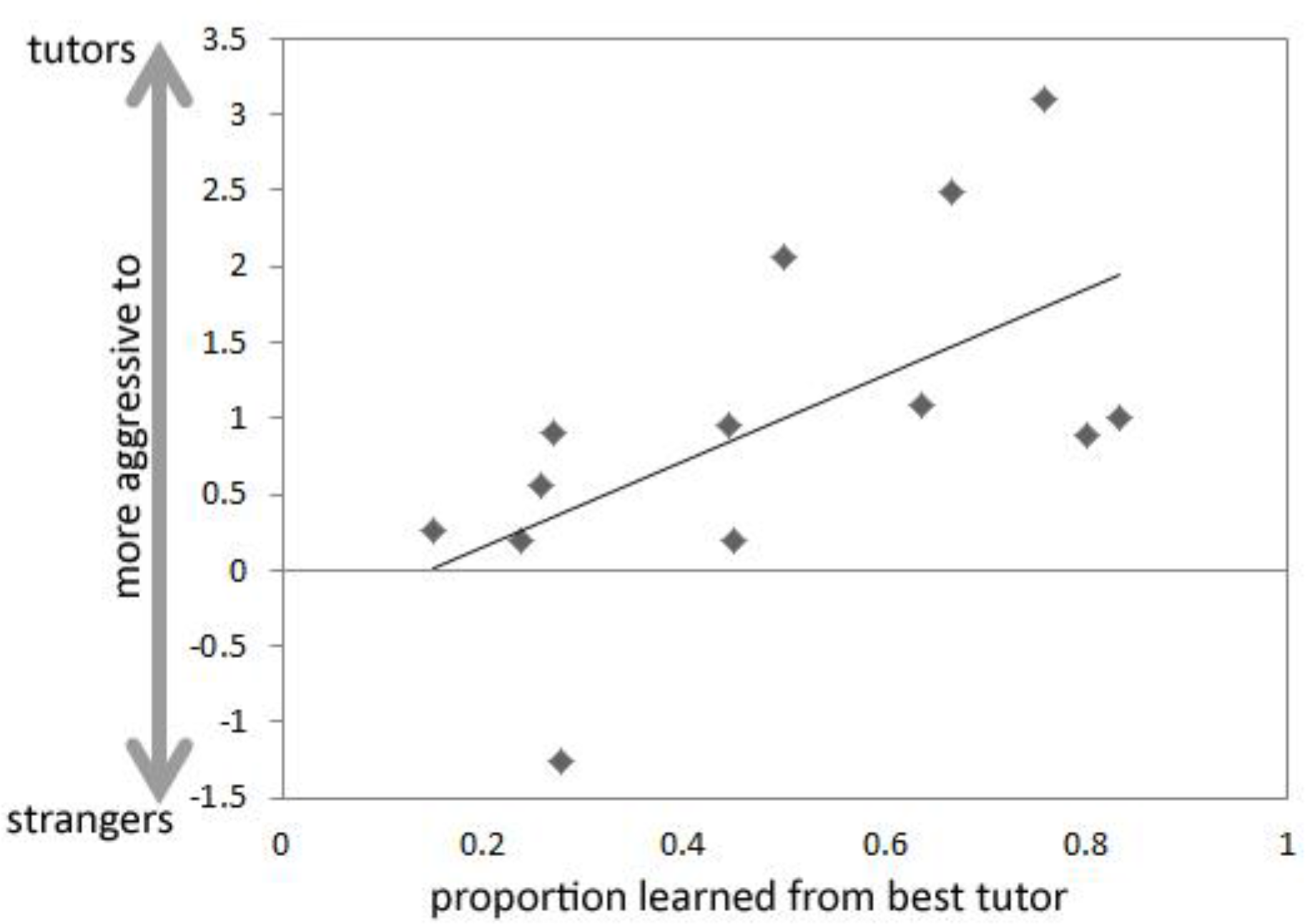
The difference in aggression scores between best tutor and stranger trials for individual subjects depending on the proportion of songs they learned from their best tutor.

## Discussion

We observed two effects in this experiment. First, subjects responded more aggressively to simulated intrusions by their former best tutors than to those of strangers. Second, the difference in response strength to tutors vs. stranger was larger the more songs the subject had learned from that tutor. These results support the competition hypothesis which predicted that tutors will elicit a higher response than strangers and that the strength of response will depend on the degree of song learning from that tutor. Below, we first discuss and critically evaluate some alternative explanations before discussing the implications of these results for the role of social interactions during song learning.

### Ruling out alternative hypotheses

Previous studies by our group and others with song sparrows in both western and eastern populations put the present findings in a fuller context and rule out certain alternative interpretations. One possible interpretation of the first effect is that birds respond more strongly to shared song than to unshared song. However, two previous studies found that song sparrows did not respond more aggressively to own (self) song, which by definition is shared compared to stranger (i.e. non-shared) song (McArthur, 1986; Searcy, McArthur, Peters, & Marler, 1981). In a more direct test, we also failed to detect differences in responses to shared vs. non-shared stranger songs (Akcay, McKune, Campbell and Beecher, in preparation). These results show that song sparrows do not in general respond more aggressively to shared songs compared to non-shared songs, eliminating this alternative explanation.

A second alternative explanation for a stronger response to tutors compared to strangers is that birds respond more strongly to local songs from tutors that used to hold territories close to subjects compared to stranger songs coming from birds that lived farther away. However, such discrimination is typically seen only over much larger distances than those involved in our study: in Searcy and colleagues’ study of eastern song sparrows (Searcy, Nowicki, Hughes, & Peters, 2002), discrimination was achieved only for non-local songs from 540 km away.

A final alternative explanation is that the present results may be explained by retaliation to a familiar neighbor (i.e. the tutors in our experiment) who broke off the Dear Enemy relationship. In our previous research we did indeed show that song sparrows increase aggression towards defecting neighbors intruding on neighbor territories (Akçay et al., 2010; Akçay et al., 2009). This retaliation however, cannot explain the present results as even a retaliation strategy does not predict a higher response to a defecting neighbor than stranger. Indeed, there have been at least five different studies by five different research groups examining responses of male song sparrows neighbors vs. strangers inside the territory of a focal bird (Harris & Lemon, 1976; Kroodsma, 1976; Moser-Purdy, MacDougall-Shackleton, & Mennill, 2017; Searcy et al., 1981; Stoddard, Beecher, Horning, & Campbell, 1991). As summarized in Table 3, the relative response strength to neighbors vs. strangers varies depending on the location, with the differences between neighbor and stranger conditions being most pronounced at the boundary with the neighbor being tested, and getting weaker or disappearing altogether at the territory center. In none of the studies however, did neighbor (or familiar) song elicit a higher response than stranger song at the territory center (see also Falls & Brooks, 1975; Stoddard, 1996). For instance, in the study most comparable to the present one, Stoddard and colleagues (1991) showed that while song sparrows are less aggressive to their neighbors when these are simulated singing at the appropriate territory boundary compared to strangers (the quintessential Dear Enemy effect), they responded equally aggressively to neighbors and strangers when the playbacks were carried out at the center of the territory (as in the present experiment) or at an inappropriate boundary (Stoddard et al., 1991; see also Stoddard, Beecher, Horning, & Willis, 1990). Overall, the conclusion from these studies suggest that song sparrows (and other songbirds) respond either equally strongly to strangers and neighbors or less strongly to neighbors than to strangers. Therefore, retaliation against a familiar individual cannot explain the present results.

**Table 3.**
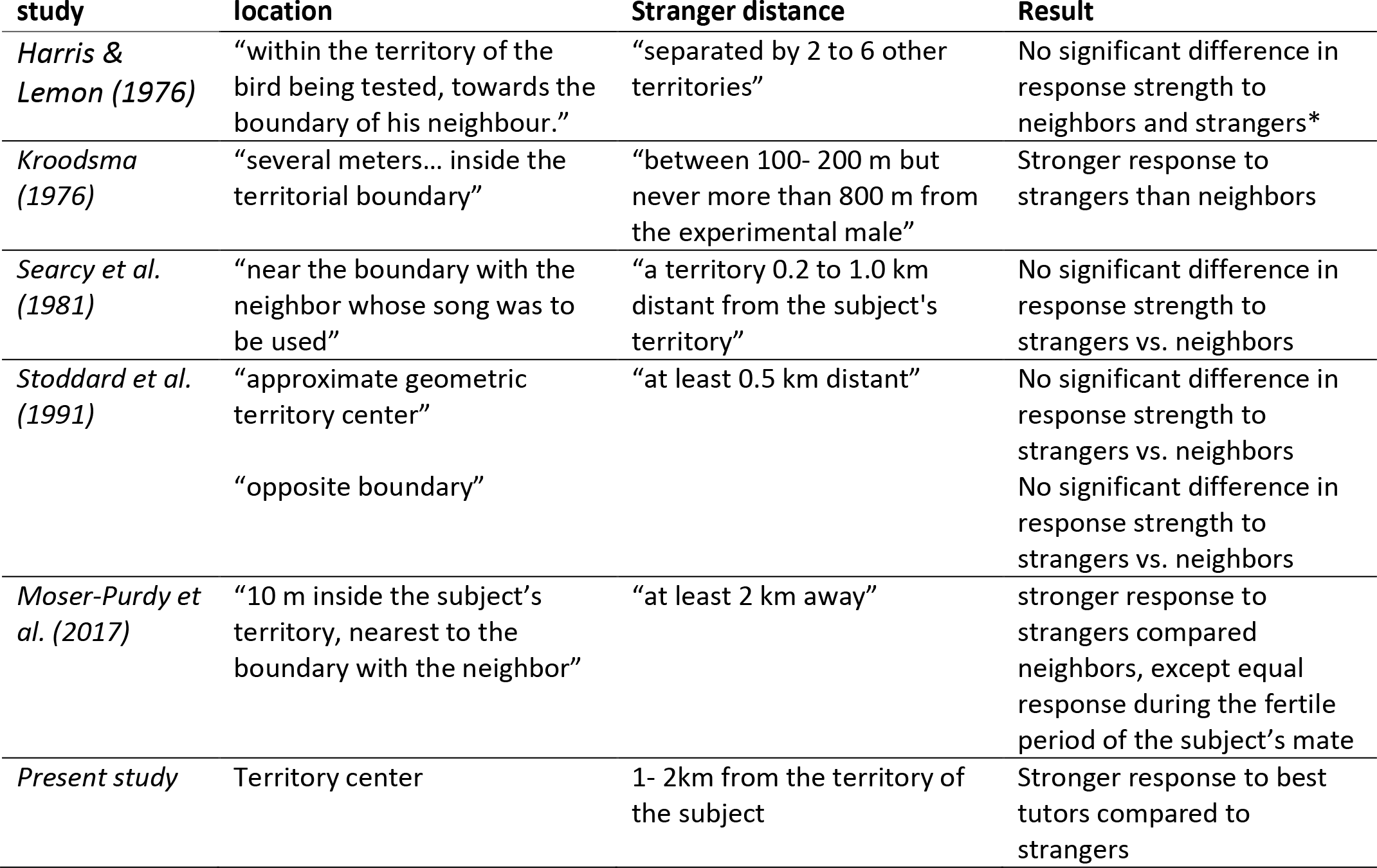
Studies with song sparrows contrasting responses to neighboring and non-neighboring (stranger) males’ song inside the territory boundaries. In each of the five previous studies stranger playbacks elicited either higher or equal responses compared to neighbor playbacks.

Given the context of these previous studies, our finding that subjects responded more strongly to tutor song than stranger song, with this difference being larger the more they had learned from the tutor, suggests that neighbors regarded the intrusion by the tutor-neighbor as a higher threat than even that by a stranger. We interpret these results as implying that the relationship between a tutor and tutee is not in a cooperative relationship. Instead, the increased aggression towards tutors point to a highly competitive relationship. These results are the first field study to indicate that birds recognize their former best tutors and that their learning history is reflected in their aggressive response to these tutors.

Note that there is a third hypothesis that might be confused with the Competition hypothesis but is in fact distinct from it. We call it simply the *Aggression hypothesis*, which states that the young birds learn the most songs from birds that are most aggressive in general. In some cases, it is further assumed that birds who are more aggressive are superior in quality to those birds who are less aggressive. We have done two previous studies that found no support for this hypothesis. In the first study (Akçay, Campbell, & Beecher, 2014; Akçay, Campbell, Reed, et al., 2014) we measured aggressiveness in adult birds in our study population and found it highly repeatable but unable to explain any of the variance in the ‘tutoring success’ of these birds in one year: young song sparrows did not learn more songs from tutors who were generally more aggressive. In the second study (Akçay et al., 2015), we compared the survival rates of birds who varied in aggressiveness and found that on average more aggressive birds do not survive longer on territory than do less aggressive birds. To be clear, the Competition hypothesis refers to a background of aggressive interactions between a specific tutor and a specific tutee, while the Aggression hypothesis refers specifically to the general effects on song tutoring of the consistent individual differences in aggressiveness (across time and across contexts) of between tutors.

### Implications for the function of song learning

One interpretation of the finding that the aggressive response to tutors co-varied positively with the amount of learning from that tutor is that young males learn more from tutors with whom they engage in more aggressive interactions. This can be considered an adaptive learning strategy given what we know about the function of song in aggressive interactions in song sparrows. Extensive field studies in our population have revealed that shared songs with neighbors are used in a graded and hierarchical signaling system (Beecher, Campbell, Burt, Hill, & Nordby, 2000; Burt et al., 2001; Searcy, Akçay, Nowicki, & Beecher, 2014). In particular, shared songs are used in two ways: as a ‘song type match’ to a neighbor singing the same song type (Beecher et al., 2000) and as a ‘repertoire match’ to a neighbor singing a different but still shared song type (Beecher, Stoddard, Campbell, & Horning, 1996). These signals indicate different levels of aggressive intention, with type matching being a reliable signal indicating willingness to escalate and eventually attack (Akçay et al., 2013; Burt et al., 2001) and repertoire matching being an intermediate signal indicating attention to the opponent’s singing but not a direct escalation. Non-shared songs are used to indicate unwillingness to continue the interaction (Beecher & Campbell, 2005). These graded signals are used in a hierarchical way such that type matching is followed by higher level threat signals such as soft songs and wing waves, and eventually physical attack if the opponent does not back down, as he could by switching to a different song type (Akçay et al., 2013; Burt et al., 2001).

Given these functions of shared songs in aggressive interaction, it is likely adaptive for young males to maximize their repertoire overlap with the adults they most often interact with aggressively. Such a strategy would allow them to mediate aggressive interactions using shared song and potentially avoid getting into physical aggression that could be costly to both parties. On the flip-side, if birds interact with multiple neighbors aggressively throughout song learning, the birds may try to overlap their song repertoire with multiple tutors by learning one or two songs from each.

A significant caveat to the present results is that they only indirectly support for the Competition hypothesis since we did not track every interaction between tutees and potential tutors during song learning. Ideally, we would want to observe the direct aggressive interactions between the tutors and tutees during the period of song learning, although previous attempts by our group using extensive radio-tracking failed to yield significant amounts of aggressive interactions between young birds and potential tutors (Templeton, Reed, Campbell, & Beecher, 2012). Nevertheless, detailed field studies have shown that new birds often do engage in repeated aggressive interactions with territory owners in order to carve out their own territory (Arcese, 1989; Nice, 1943). It is possible that intense aggressive interactions mostly happen in a limited time frame when the young bird first establishes his territory, which in our population can happen any time between their first Summer (as early as July and August) and the following Spring (as late as May). More detailed studies are needed, particularly ones that would take advantage of automated tracking systems that can monitor tutors and tutees around the clock for extended periods of time (e.g. Rutz et al., 2012). Such automated systems could be used to detect when territories are first established and to quantify how many interactions the young birds have with their neighbors during this period. Until then, the present results provide only tentative support of the competition hypothesis.

